# A non-coding genetic variant maximally associated with serum urate levels is functionally linked to HNF4A-dependent PDZK1 expression

**DOI:** 10.1101/362277

**Authors:** Sarada Ketharnathan, Megan Leask, James Boocock, Amanda J. Phipps-Green, Jisha Antony, Justin M. O’Sullivan, Tony R. Merriman, Julia A. Horsfield

## Abstract

Several dozen genetic variants associate with serum urate levels, but the precise molecular mechanisms by which they affect serum urate are unknown. Here we tested for functional linkage of the maximally-associated genetic variant *rs1967017* at the *PDZK1* locus to elevated *PDZK1* expression.

We performed expression quantitative trait locus (eQTL) and likelihood analyses followed by gene expression assays. Zebrafish were used to determine the ability of *rs1967017* to direct tissue-specific gene expression. Luciferase assays in HEK293 and HepG2 cells measured the effect of *rs1967017* on transcription amplitude.

PAINTOR analysis revealed *rs1967017* as most likely to be causal and *rs1967017* was an eQTL for *PDZK1* in the intestine. The region harboring *rs1967017* was capable of directly driving green fluorescent protein expression in the kidney, liver and intestine of zebrafish embryos, consistent with a conserved ability to confer tissue-specific expression. The urate-increasing T-allele of *rs1967017* strengthens a binding site for the transcription factor HNF4A. siRNA depletion of HNF4A reduced endogenous *PDZK1* expression in HepG2 cells. Luciferase assays showed that the T-allele of *rs1967017* gains enhancer activity relative to the urate-decreasing C-allele, with T-allele enhancer activity abrogated by HNF4A depletion. HNF4A physically binds the *rs1967017* region, suggesting direct transcriptional regulation of *PDZK1* by HNF4A.

With other reports our data predict that the urate-raising T-allele of *rs1967017* enhances HNF4A binding to the PDZK1 promoter, thereby increasing *PDZK1* expression. As PDZK1 is a scaffold protein for many ion channel transporters, increased expression can be predicted to increase activity of urate transporters and alter excretion of urate.

## INTRODUCTION

Hyperuricemia is necessary but not sufficient for gout (1). Serum urate levels have a considerable heritable component (2,3). Therefore understanding the molecular mechanisms of serum urate control will reveal important knowledge on the etiology of gout. Genome-wide association studies (GWAS) in several populations to detect genetic variants controlling serum urate levels have identified several dozen loci (4-7). Most of these loci also associate with gout (4,8,9). The loci of strongest effect include genes encoding renal and gut uric acid transporters (e.g. SLC2A9, ABCG2, SLC22A12) and their accessory molecules (e.g. PDZK1). The well-characterized ABCG2 p.Gln141Lys protein variant is highly likely to be causal (10). However, the causal genes and genetic variants at the considerable majority of the serum urate loci remain unconfirmed (11). Co-localization analysis of GWAS and expression quantitative trait loci (eQTL) has identified instances where genetic variation correlates with changes in gene expression and this strategy has identified 19 candidate causal genes for serum urate control (7).

The PDZ (PSD-95, DglA, and ZO-1) domain-containing protein PDZK1 is a binding partner of the canonical uric acid urate transporter 1 (URAT1). PDZK1 also physically interacts, and has co-ordinate expression with, the majority of transporters for uric acid (12). For example in human kidney samples PDZK1 protein and mRNA levels correlate with a range of apical transporters (13) and PDZK1 has been implicated in regulation of the expression of ABCG2 in human intestinal cell lines (14). Protein-protein interaction occurs via contact between PDZK1 and a PDZ-interacting domain present at the extreme C-terminal end of the various transporters. This has led to the idea that PDZK1 acts as a scaffold for a uric acid ‘transportosome’ (15); a function that implicates PDZK1 as a key molecule in urate homeostasis (12). PDZK1 has the potential to regulate the cellular location of, and/or stabilize, uric acid transporters and co-transport molecules. Therefore the regulation of PDZK1 levels would be expected to contribute to urate homeostasis, making understanding of the molecular mechanisms that control PDZK1 expression important. This is particularly important in the context of reports of association of PDZK1 with gout in multiple ancestral groups (8,16,17) and the established role of renal and gut under-excretion of uric acid as a cause of gout (18,19). Given that a urate GWAS association signal maps to a small region (<10 kb) immediately upstream of the transcription start site of *PDZK1* (4,20), our aim was to use *in vivo* and *in vitro* approaches to understand if variants in proximity to *PDZK1* control serum urate levels by modulating transcription of this gene.

## RESULTS

### Association with serum urate levels and *PDZK1* expression – *rs1967017* as a candidate causal variant

We used imputation to update the Kottgen *et al*. analysis performed in 2012 (4) using the latest reference haplotypes (figure 1A). This revealed the maximal region of association to reside in a tightly-defined 1 kb region (145.711 Mb) upstream of the *PDZK1* transcription start site. Probabilistic Annotation Integrator (PAINTOR) analysis (21) suggested that of all the associated SNPs, *rs1967017* is the most likely to be causal (Z-score 11.2, PAINTOR *p* = 0.93) (Table S1). The *rs1967017* SNP is located 4 kb upstream of the transcription start site of the *PDZK1* gene. Analysis of the GTEx database (22) revealed that *rs1967017* is a strong eQTL for PDZK1 in colon (*p* = 1.3 x 10^-11^) and small intestine (*p* = 8.3 x 10^-7^) (figure S1A). *rs1967017* is a weaker eQTL for PDZK1 in the pancreas and there was no evidence for an eQTL in the liver or stomach. There are currently no renal eQTL data available in GTEx. However three separate studies provide no evidence for *rs1967017* as an eQTL in the kidney (23,24) or in the glomerulus (25 figure S1B). Co-localisation analysis using COLOC (26) gave a posterior probability of 0.93 for the serum urate GWAS and *PDZK1* colon eQTL signals to co-localise (figure 1B). The similarity between the signals is also evident from figure 1. Using the tightly-associated surrogate marker *rs1471633* (LD with *rs1967017* is r^2^=1 in European), we determined that the urate-increasing allele (A for *rs1471633*, completely correlated with T for *rs1967017*) associated with increased expression of *PDZK1*, with reduced expression requiring two copies of the urate-decreasing allele (C for *rs1471633*, completely correlated with C for *rs1967017*) in the colon, with only one copy required in the small intestine (figure S1A).

**Figure 1.**
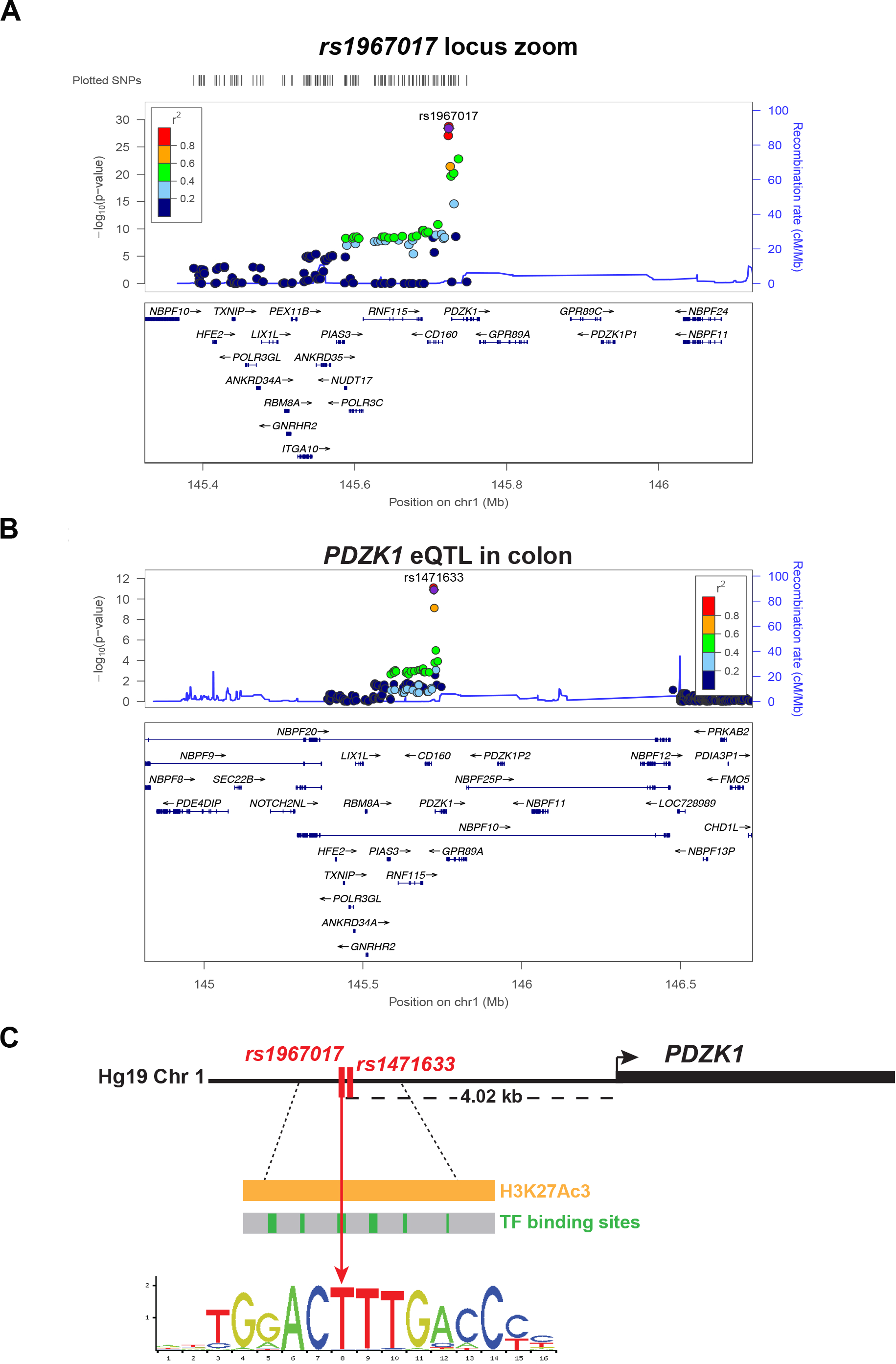
In silico analysis of the urate-associated GWAS SNP *rs1967017* (C>T). **(A)** A regional association plot for serum-urate associated GWAS SNPs (4) present in close proximity to the *PDZK1* gene. Each SNP is colored based on its correlation with the lead SNP, *rs1967017*. **(B)** Cis eQTL for *PDZK1* in colon. The figure was generated using LocusZoom. Publicly-available GTEx data were used from the Database of Genotype and Phenotype (www.ncbi.nlm.nih.gov/gap) under project #834. **(C)** *rs1967017* maps 4.02kb upstream of the *PDZK1* gene in a region rich in enhancer histone modifications (orange bar) and transcription factor binding sites (green boxes) (ENCODE). Presence of the serum urate-raising *rs1967017* T allele results in an HNF4A site with increased binding affinity, as predicted by JASPAR. The second maximally-associated SNP at the locus (*rs1471633*) is located 94 bp away from, and is in complete linkage disequilibrium with, *rs1967017*.

Using ENCODE (27), we determined that the region harboring *rs1967017* is contained within a transcriptionally-active domain (H3K27ac3, H3K4me1, H3K4me2; figure S2). Analysis of transcription factor binding in multiple cell lines showed that several transcription factor binding sites are present in the region, including those recruiting HNF4A, although HNF4A binding is restricted to the HepG2 cell line. Interestingly, there is a signal of association of serum urate at the *HNF4A* locus in the Kottgen *et al*. (4) GWAS (figure S3). Significantly, the urate-raising T allele of *rs1967017* that is associated with increased expression of *PDZK1* markedly strengthens an existing HNF4A binding site, with the binding score increasing 4-fold (from 2.645 to 8.845) when the T allele is present, compared with the C allele (figure 1C) (28) Together, the data indicate that *rs1967017* resides in a region that controls *PDZK1* expression and that the T>C variant is likely to affect *PDZK1* transcription.

### The *rs1967017* regulatory region drives gene expression in kidney, liver and intestine

Conservation of key functional genes (such as those involved in urate transport) through evolution enables the use of animal models to clarify spatiotemporal gene expression. We used the zebrafish animal model as a read-out to determine in which tissues the *rs1967017* regulatory region might drive gene expression. Human *PDZK1* is expressed at the highest level in the kidney, and is also expressed in the liver and intestine (Table 1). Zebrafish *pdzk1* is expressed in kidney, nephric duct and intestinal bulb at 5 days post-fertilization (dpf) (29). The similar expression profiles of the human and fish *Pdzk1* orthologs supports the hypothesis that regulatory information driving expression of *PDZK1/pdzk1* is conserved.

**Table 1.** GTEx gene expression data for *PDZK1* and *HNF4A*.

| **Tissue** | ***PDZK1* expression** | ***HNF4A* expression** |
| --- | --- | --- |
| **Kidney cortex** | 55.495 | 8.149 |
| **Small Intestine** | 20.597 | 28.229 |
| **Liver** | 18.777 | 39.978 |
| **Colon (transverse)** | 0.269 | 29.992 |
| **Pancreas** | 2.815 | 4.523 |
| **Stomach** | 0.255 | 4.356 |

Values shown are median RPKM (Reads Per Kilobase of transcript per Million mapped reads).

We created stable transgenic zebrafish lines using an 857 bp putative regulatory region containing either allele of *rs1967017* (C or T) fused to enhanced green fluorescent protein (eGFP). GFP expression was observed as early as 13-14 hours post-fertilization (hpf; 12-somite stage, data not shown). In 4 days post-fertilization (dpf) transgenic zebrafish embryos, expression driven by the T allele (figure 2A-A’’) and the C allele (figure 2B-B”) was observed in the glomerulus (red arrows), the nephric ducts (orange arrows) and the intestinal epithelium (yellow arrows). Confocal analysis at 5 dpf confirmed expression in the same locations (figure 2C, D). Expression of eGFP in the *rs1967017*-C/T-GFP transgenic lines was very similar to the endogenous expression pattern of *pdzk1* expression in zebrafish (29), and consistent with the expression of *PDZK1* in human (Table 1). The expression pattern driven by *rs1967017* C allele (urate decreasing in humans) and *rs1967017* T allele (urate increasing in humans) was spatiotemporally identical. However, expression driven by the T allele was noticeably less robust in the glomerulus than that driven by the C allele (figure S4), indicating that the T>C base change confers subtle differential activity in zebrafish. To determine more precisely the identity of tissue expression in zebrafish, we sectioned 21-day-old larvae and analyzed *GFP* mRNA distribution by *in situ* hybridization. We found intense GFP staining in the liver and in the intestinal epithelium of zebrafish, with less intense staining present in the kidney (figure 2E). This distribution reflects the relative levels of *HNF4A* expression in human tissues from GTEx (Table 1). GFP expression is also coincident with the previously reported expression of *hnf4a* in zebrafish kidney, liver and intestine (30,31).

**Figure 2.**
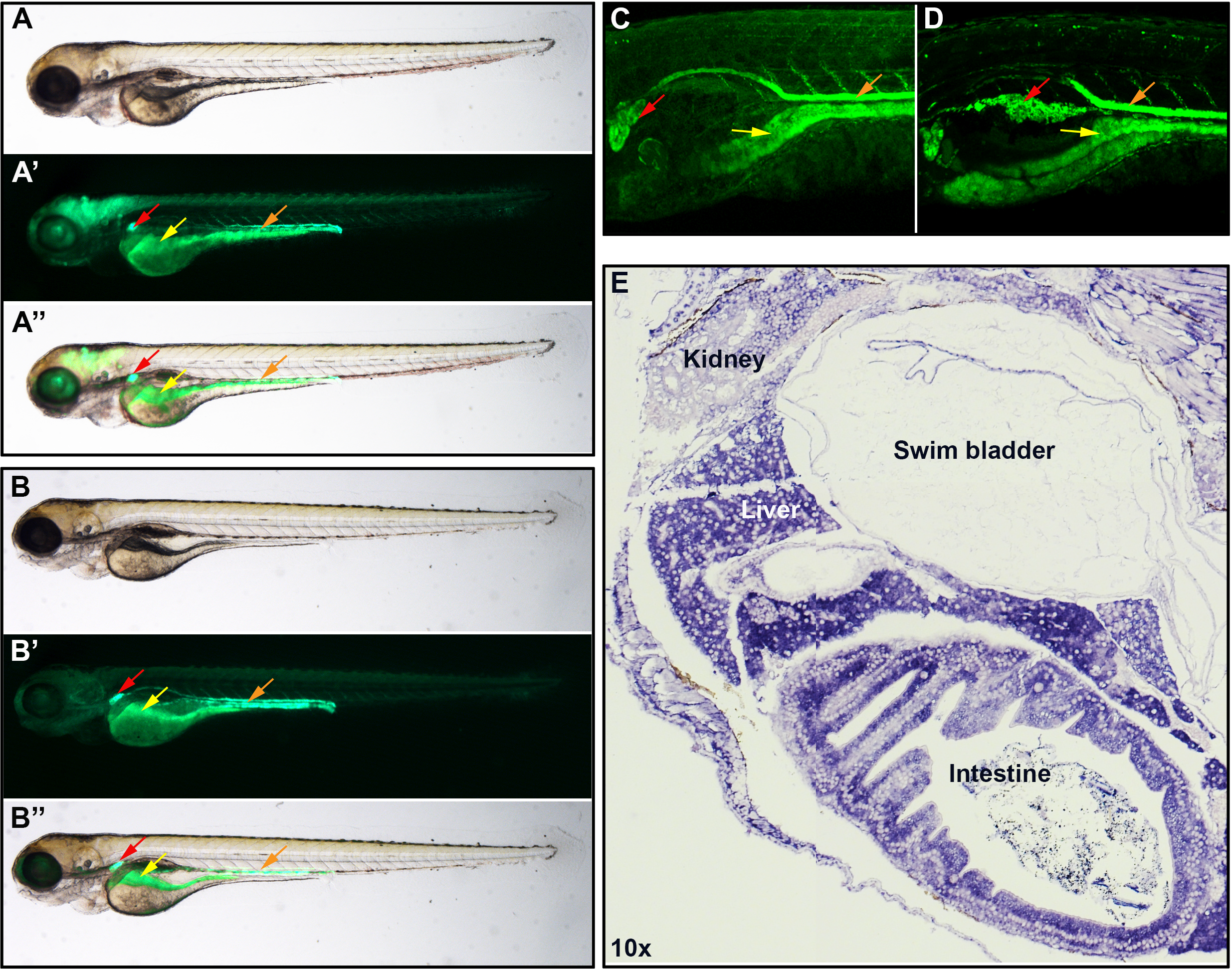
*rs1967017* putative regulatory human DNA recapitulates *pdzk1* expression in zebrafish. Whole mount representative **(A, B)** lateral bright field, **(A’, B’)** GFP fluorescence and **(A’’, B’’)** merged images of 96 – 120 hpf F2 zebrafish embryos carrying **(A, A’, A’’)** human *rs1967017* protective ‘C’ allele-GFP DNA and **(B, B’, B’’)** urate-raising T’ allele-GFP DNA. **(C)** Confocal sections of 120 hpf (4-5 dpf) F2 zebrafish embryos carrying the human *rs1967017* urate-lowering C allele-GFP and **(D)** urate-raising T allele-GFP. Both alleles drive expression in the glomerulus (red arrows), pronephric duct (orange arrows) and intestinal epithelium (yellow arrows). **(E)** A sagittal section through the midline of a 21 dpf zebrafish carrying the human *rs1967017* urate-lowering C allele-GFP was subjected to in situ hybridization with a riboprobe detecting *GFP* mRNA. Robust GFP expression can be seen in the kidney, liver and intestine.

The results indicate that the human DNA sequence harboring the *rs1967017* SNP has sufficient information to recapitulate the normal expression pattern of *PDZK1*. Moreover, although information conferring tissue identity was not affected by the variant base in the *rs1967017* SNP, the C>T base variation affects quantity and/or stability of expression in zebrafish glomerulus (figure s4). Together, the results suggest it is unlikely that *rs1967017* affects serum urate via altered spatiotemporal expression of *PDZK1*.

### The *rs1967017* urate-raising T allele increases gene expression in human cells

To determine if the regulatory region containing *rs1967017* influences gene expression in human cells, we cloned the 857 bp region (described above) into a plasmid expressing luciferase (pGL4.23). Using a dual luciferase reporter system, we determined that this region acts as an enhancer in two different cell lines, HEK293 (human embryonic kidney) and HepG2 (liver) (figure 3). In HEK293 cells, both alleles (C and T) were able to enhance luciferase expression (figure 3A). In HepG2 liver cells, the urate-decreasing C allele caused a small but non-significant increase in expression. In contrast, the urate-increasing T allele was highly active in HepG2 liver cells, conferring ~6-fold more luciferase expression than it did in HEK293 cells (figure 3B). In both cell lines, the T allele was more active than the C allele, with activity increased over the C allele by ~2-fold in HEK293 and ~10-fold in HepG2.

**Figure 3.**
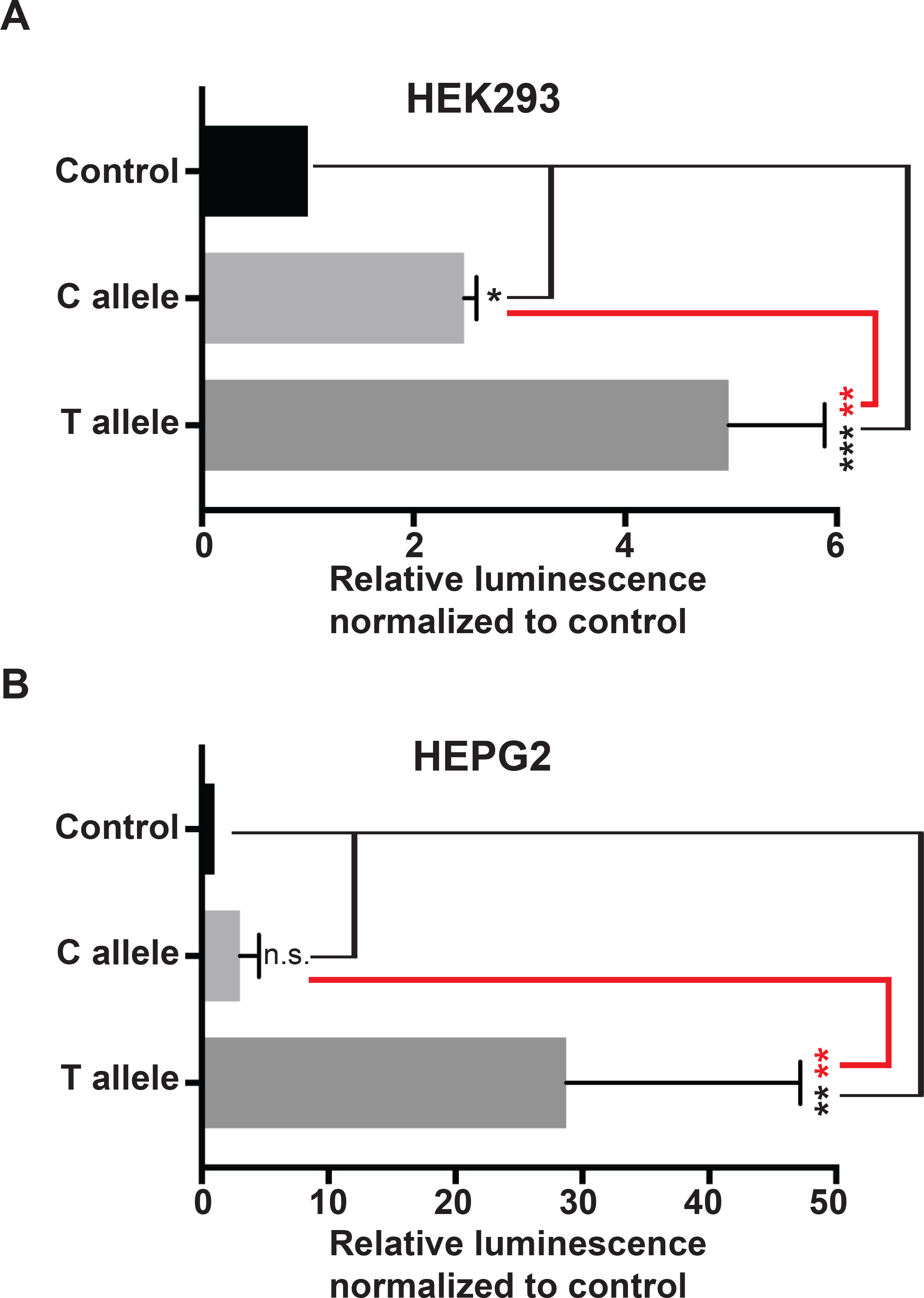
The *rs1967017* T allele enhances gene expression in renal and hepatic cell lines. Enhancer activities of pGL4.23 (control), pGL4.23 containing the *rs1967017* urate-lowering C allele-GFP DNA and pGL4.23 containing the *rs1967017* urate-raising T allele-GFP DNA were measured using a dual luciferase system. **(A)** Both alleles show significant enhancer activity in HEK293 cells. **(B)** Only the urate-raising T allele is a significant enhancer in HepG2 cells. The average of at least three biological replicates was calculated and statistical significance determined using Tukey’s multiple comparisons test. Error bars denote standard deviation and asterisks indicate significance: * p<0.05; ** p<0.005; *** p<0.0005; NS – not significant; red asterisks – C allele versus T allele.

### The *rs1967017* urate-raising allele is dependent on HNF4A for enhancing transcription in HepG2 cells

We investigated whether the urate-increasing T allele, which strengthens an HNF4A binding site (figure 1), confers increased transcriptional activity mediated by HNF4A expression. Notably, the kidney cell line HEK293 has very low mRNA levels of HNF4A in contrast to HepG2 where it is robustly expressed (figure S5, S6). Because the T allele is a strong enhancer of luciferase expression in HepG2, we were interested to determine if knock-down of HNF4A in HepG2 cells would block the ability of the T allele to increase luciferase transcription (figure 4).

**Figure 4.**
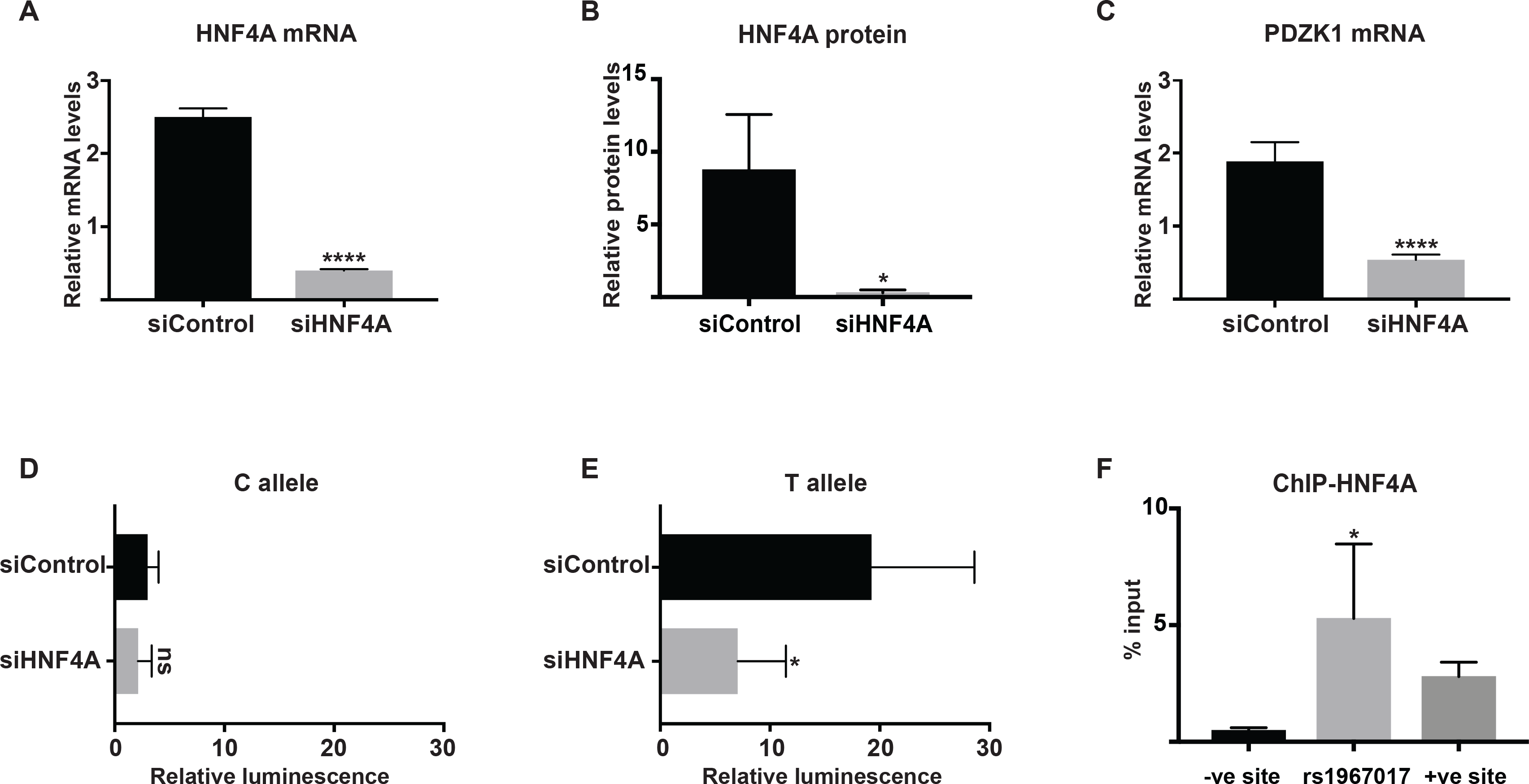
*rs1967017* recruits HNF4A to directly regulate *PDZK1* expression in HepG2 cells. HepG2 cells were reverse-transfected with either 10 nM *HNF4A* siRNA or 10 nM of a control siRNA. Cells were collected and analysed 48 hours post-transfection. **(A)** *HNF4A* mRNA expression is reduced, **(B)** HNF4A protein expression is abolished, and **(C)** endogenous *PDZK1* mRNA is downregulated in *HNF4A* siRNA-treated cells. Expression of *GAPDH* and *PPIA* mRNA were used for the normalization of mRNA expression and GAPDH protein levels for normalization of protein expression. **(D, E)** Enhancer activity of the *rs1967017* urate-lowering C allele (D) and urate-raising T allele (E) in *HNF4A* siRNA-treated cells versus control HepG2 cells. Results represent the average of five biological replicates, error bars are ± S.D., and statistical significance was determined using an unpaired *t*-test. **(F)** Quantitative PCR detection of HNF4A binding to the *rs1967017* locus at the PDZK1 promoter by chromatin immunoprecipitation using an anti-HNF4A antibody. Negative and positive binding sites for HNF4A were promoter regions of β-globin and ABCC6 (31), respectively. Results represent the average of three biological replicates, error bars are ± S.D. Statistical significance determined using Tukey’s multiple comparisons test. and asterisks indicate significance: * p<0.05; *** p<0.0005; **** p<0.00005.

siRNA knockdown of HNF4A in HepG2 cells was effective in reducing HNF4A mRNA (figure 4A) and protein (figures 4B, S6) levels. We also observed down regulation of endogenous *PDZK1* mRNA upon depletion of HNF4A (figure 4C). Depletion of HNF4A resulted in a ~3-fold reduction in the ability of the T allele to enhance luciferase expression, whereas expression driven by the C allele was not affected by HNF4A reduction (figure 4D, E). Therefore, in HEPG2 cells that express HNF4A, the enhancer activity of the T allele is dependent on maintenance of HNF4A levels.

### HNF4A binds to the *rs1967017* region in HepG2 cells

To determine if HNF4A directly regulates the *rs1967017* enhancer region, we performed a quantitative ChIP assay in HepG2 cells, which are heterozygous for *rs1967017* (32). Following HNF4A immunoprecipitation, a 202 bp fragment was amplified using primers flanking the target HNF4A motif. An *ABCC6* promoter region bound by HNF4A (33) was used as positive control, and human β-globin as negative control. Quantitative amplification of HNF4A at the *rs1967017* region was approximately double the levels of the ABCC6 positive control, confirming that HNF4A binds the *rs1967017* enhancer region (figure 4F). To determine if the urate-increasing T allele is bound preferentially to the urate-decreasing C allele, we deep-sequenced the 202 bp immunoprecipitated amplicon containing the target motif in samples before and after HNF4Aimmunoprecipitation. From this experiment, we were not able to conclusively determine a preference of HNF4A for the T allele, although there was a trend for preferential immunoprecipitation of the T allele in one of two replicates (Table S2).

## DISCUSSION

Many genetic variants that contribute to the risk of complex phenotypes act by influencing expression of genes, rather than by altering their coding sequence (34). Here, we show that one of the maximally associated SNPs at *PDZK1* (*rs1967017*) and most likely to be causal by PAINTOR analysis directly affects *PDZK1* expression levels. A DNA segment containing *rs1967017* was able to drive gene expression in zebrafish and in two human cell lines. The 857 bp DNA segment had sufficient information to confer kidney, liver and intestine expression in zebrafish, matching the tissue expression profile of *PDZK1* in human, suggesting that regulation of this gene is conserved through evolution. Interestingly, we observed expression driven by the *rs1967017* enhancer in the epithelial cells of the zebrafish intestine, which is a cell type in which one would expect to find solute transporters. While the identity of the polymorphic variant nucleotide at *rs1967017* generally did not affect tissue-specific gene expression in zebrafish, it did modulate the level of expression observed in the kidney, with the urate-increasing T allele conferring weaker expression than the C allele. This is consistent with our understanding of the composite nature of gene regulation and likely reflects the presence of a different complement (or activity) of transcription factors in kidney, compared with gut, available to enhance transcription from the *rs1967017* element.

There was a robust and reproducible effect of the variant nucleotide at *rs1967017* on level of gene expression in two human cell lines. HEK293 is a cell line frequently used to model human kidney (35) and in this line, the urate-increasing T allele enhanced gene expression comparative to the C allele. However, the T allele was 10-fold more active in the liver cell line HepG2. The reason for this is likely to be that the T variant strengthens a consensus binding sequence for HNF4A, a transcription factor that is expressed in HepG2 but not HEK293. Supporting a causal role for *rs1967017*, *PDKZ1* expression is activated by the nuclear receptor thyroid hormone receptor β (THRB) in HEPG2 cells, with the T allele of *rs1967017* associated with increased transactivation by THRB compared to the C allele (24). This is not observed for either allele of *rs1471633*, the other maximally urate-associated variant at the locus (figure 1). In our data the enhanced reporter expression was substantially reduced when HNF4A was knocked down in HepG2 cells, and was accompanied by down regulation of endogenous *PDZK1*. Our results are consistent with the report that the zebrafish *pdzk1* promoter is bound by *hnf4a* and that *pdzk1* expression in the zebrafish digestive tract is *hnf4a* dependent (31). Furthermore, regulation of *PDZK1* by HNF4A is likely to be direct, since HNF4A physically binds the *rs1967017* element (figure 4F). Although we could only obtain suggestive preliminary evidence that the T allele is bound preferentially bound by HNF4A over the C allele, the eQTLs of *rs1967017* with PDZK1 and the fact that variants at HNF4A also associate with serum urate levels give confidence to the hypothesis that HNF4A modulates *PDZK1* expression as part of an enhancer complex located within the locus marked by *rs1967017*.

PDZK1 is a scaffolding protein involved in assembly of a transporter complex in the apical membrane of the proximal tubule of the kidney (12). Given that the urate-increasing T allele at *rs1967017* associates with increased expression of *PDZK1*, it is reasonable to assume that it increases PDZK1 protein levels. The importance of the ratio of PDZK1 to uric acid transport proteins (such as URAT1, encoded by SLC22A12) that it interacts with in the kidney is unknown, but it is established that PDZK1 stabilizes transporter localization in the membrane and increases transporter activity (12,15). It was interesting to note that only tissues with very strong evidence for an eQTL were the colon and small intestine, with no evidence for an eQTL in the kidney (23-25). It is possible that other regulatory mechanisms exist in the kidney instead of the *rs1967017*-mediated regulatory mechanism of *PDZK1* expression observed in the gut. For example it has recently been shown that HNF1A positively regulates *PDZK1* expression and binds to the *PDZK1* promoter in the human kidney (13). In figure 5, we provide a model for the mechanism of action of the *rs1967017* variant that best explains the collective data. We propose that the enhancer variant *rs1967017* modulates PDZK1 expression in the intestine (an important site of urate excretion) but not in the kidney. *Rs1967017*-mediated regulation of expression PDZK1 in intestine would depend on the transcription factor HNF4A, which is highly expressed in intestine and liver relative to kidney (www.gtexportal.org), and which physically binds to the enhancer containing *rs1967017*. The T-allele of *rs1967017* strengthens the binding site for HNF4A and increases the expression of PDKZ1 in the gut. Increased HNF4A binding leads to increased PDZK1 protein, which is predicted to stabilise the transportosome and increase the activity of uric acid transporters, such as ABCG2 (14,36). In human kidney samples there is no association between *rs1967017* and *PDZK1* expression (23-25). Therefore we propose that HNF1A-dependent regulation of PDZK1 expression independent of *rs1967017* is the dominant mechanism in the kidney.

**Figure 5.**
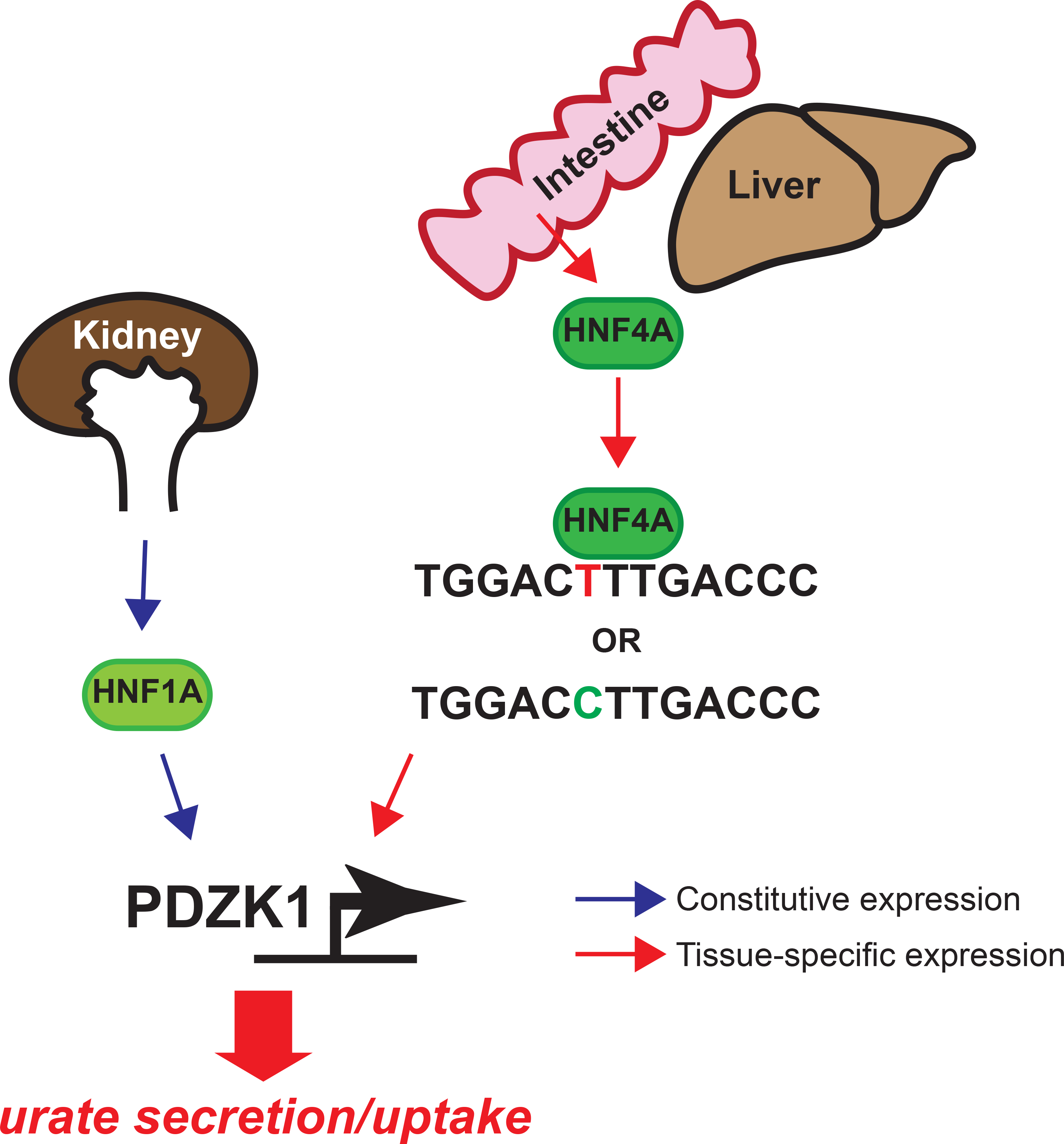
Model for molecular mechanism of *rs1967017*. The serum urate variant *rs1967017* lies ~4 kb upstream of the gene encoding ion channel scaffold protein, PDZK1, in a regulatory element that controls gene expression. PDZK1 is constitutively expressed in kidney under the control of HNF1A (13) and other factors. In contrast, the enhancer variant *rs1967017* modulates PDZK1 expression not in the kidney, but rather, in the intestine. Regulation of PDZK1 in intestine depends on the transcription factor HNF4A, which is highly expressed in intestine and liver relative to kidney, and which physically binds to the *rs1967017* enhancer. The T-allele of *rs1967017* strengthens the binding site for HNF4A and increases its expression in the gut, where HNF4A and PDZK1 are co-expressed. Increased HNF4A binding leads to increased PDZK1 protein, which is predicted to stabilise the transportosome and modulate uptake and/or secretion of serum urate.

Approximately one-third of the excretion of uric acid occurs via the intestine, with the remainder via the kidney (12). Activity of the ABCG2 secretory uric acid transporter is very important in gut excretion (18), and PDZK1 is important in the correct localization of ABCG2 to the apical membrane in the gut (14,36). Urate is primarily generated in the liver from glycolysis and purine catabolic and recycling pathways (37). In the mouse, liver-specific knockout of SLC2A9 (that encodes GLUT9) results in hyperuricemia that results from blockage of the transport of urate into the liver for degradation by uricase (38). Humans have an inactive uricase and therefore substantially higher serum urate levels than mouse. Although SLC2A9 is expressed in the liver (39), the direction of hepatic transport of uric acid in humans is unknown. However, given that SLC2A9 has a typical PDZK1-binding PDZ motif (12) it is possible that increased hepatic expression of PDZK1 will stabilize SLC2A9 and thereby contribute to hyperuricemia, perhaps via increasing hepatic export of urate into the blood. Both the intestine and liver also express the transcriptional regulator HNF4A, and as outlined above, the urate-increasing T allele of *PDZK1 rs1967017* is likely to have a stronger effect where HNF4A is expressed. Further emphasizing the importance of HNF4A in urate control there is evidence that HNF4A is part of a transcriptional network regulating uric acid transporters (including GLUT9 (*SLC2A9*), OAT4 (*SLC22A11*), NPT1 (*SLC17A1*) and NPT4 (*SLC17A3*) (40,41)). Table 1 shows that while HNF4A is modestly expressed in the kidney cortex, its expression is higher in liver and intestine. Taken together, our results show that the urate-increasing allele of *rs1967017* strengthens a strong HNF4A binding site close to the *PDZK1* promoter, which is in turn likely to increase PDZK1 expression in cells that express HNF4A.

In summary, our results provide evidence for a mechanism for variant *rs1967017* to affect serum urate via altered *PDZK1* expression. It is likely that the urate-increasing variant increases transcription overall and effects PDZK1 levels through HNF4A.

## MATERIALS AND METHODS

### Imputation of the *PDZK1* region

The summary level urate GWAS data were downloaded from: http://metabolomics.helmholtz-muenchen.de/gugc/). Single nucleotide polymorphisms not present in the most recent 1000 Genomes release were removed, as were those where alleles were non-identical between the summary level data and 1000 Genomes. The effective sample size for each SNP was calculated using the Genome-wide Complex Trait Analysis (GCTA, v1.25.2) toolkit (42) and SNPs with effective sample sizes > 2 standard deviations from the mean were excluded. Finally, SNPs with a minor allele frequency (MAF) of less than 0.01 in the European population were excluded. ImpG (v1.0) (43) was used to impute *z*-scores into the summary statistics. For the reference haplotypes, the 1000 Genomes data set was used (44).

### Vectors and constructs

An 857 bp DNA fragment (chr1:145,723,012-145,723,868) harboring *rs1967017* was amplified from the genomic DNA of two individuals from the 1000 Genomes project (namely, HG00116 and HG00139) who were homozygous for the C and T alleles, respectively. Primers F: 5’-GGGCCACTAGGCTGTTCAT-3’ and R: 5’-CAGGTCTGATTGTCCTAGTATGC-3’ were used for amplification. Amplicons were cloned into pCR^®^8/GW/TOPO (Invitrogen) entry vector and gateway-cloned into the destination vectors pGL4.23-GW (Addgene 60323) (45) and pTol2Gata2 (46) using LR Clonase^®^ enzyme (Invitrogen) for the enhancer assays in human cell lines and zebrafish expression assays, respectively. The vector construct used in the enhancer assay has previously been tested for different test enhancer regions that give a range of tissue-specific expression (47) - the GFP-expression pattern we observed was unique for the *rs1967017*-containing fragment.

### *In silico* analysis of transcription factor binding sites

Transcription factor analysis on the 837 bp regulatory fragment was performed using three in silico tools. Using the ‘Transcription Factor ChIP-Seq track’ from ENCODE we analyzed our sequence for the presence of 161 transcription factors in 91 cell types. We found *rs1967017* to alter the binding affinity of only the transcription factor HNF4A (MA0114.3) in the HepG2 cell line. The position weight matrix probabilities for the T and C alleles were 0.90 and 0.009 respectively. We confirmed the same finding using the HaploReg v4.1 tool (48), the PWM scores were 16.1 and 13 for the T and C allele respectively. In addition, we scanned our DNA sequence containing either the T allele or the C allele, using the JASPAR CORE 2018 database, which has a total of 719 transcription factor profiles (537 human and 182 rat/mouse), with a relative profile score threshold of 75%. The predicted binding scores for the C and the T alleles were 2.7 and 8.9 respectively.

### Cell culture and transfections

Human embryonic kidney HEK293 cells and human hepatocellular carcinoma HepG2 cells were grown (37 °C, 5% CO_2_) and maintained in Dulbecco’s Modified Eagle’s Medium (DMEM) (11995-065, Gibco) and MEM α, no nucleosides media (12561049, Gibco) respectively, supplemented with 10% fetal bovine serum (Moregate, New Zealand). For the luciferase assay, HEK293 and HepG2 cells were transiently transfected with the pGL4.23 constructs and Renilla, using Lipofectamine 3000 (Invitrogen). Luminescence was measured after 48 hours using the Dual Glo Luciferase Assay (Promega), on the Perkin Elmer Victor X4 plate reader. Luciferase values were normalized to Renilla and to empty vector controls.

For the siRNA knockdown experiments, HepG2 cells were reverse-transfected with 10 nM scrambled control siRNA or 10 nM HNF4A siRNA (Dharmacon L-003406-00-0005) using RNAiMax (Invitrogen), concurrently for gene expression analysis and for luciferase assay. After 24 hours, culture media was replaced with fresh MEM α and the cells were immediately transfected with luciferase reagents. Luminescence was measured 48 hours post-transfection and was normalized to Renilla.

### Quantitative PCR

Total RNA was isolated from control siRNA- and HNF4A siRNA-treated HepG2 cells at 48 hours post-treatment using NucleoSpin RNA kit (Macherey-Nagel) and cDNA was synthesized with qScript cDNA SuperMix (Quanta Biosciences). *HNF4A* and *PDZK1* expression was measured using SYBR Premix Ex Taq II (Takara) on the Roche LightCycler400. Primers used were HNF4A-F: 5’-GCGTGGTGGACAAAGACAAG-3’, HNF4A-R: 5’ AGCTTGACCTTCGAGTGCTG-3’, PDZK1-F: 5’-AGCCCCGAATTGTGGAGATG-3’ and PDZK1-R: 5’-TTTCCACAGACTCGCCGTTG-3’. Reference genes were *GAPDH* and *PPIA*. Normalization was performed in qbase+.

### Immunoblot analysis

Total protein was isolated from control siRNA- and HNF4A siRNA-treated HepG2 cells at 48 hours post-treatment by lysis in Radio-Immunoprecipitation Assay buffer containing protease inhibitor cocktail (Complete™, Roche). Protein concentration was measured using Bicinchoninic Acid assay kit (Pierce, Thermoscientific). 30 µg protein per sample was separated on a 10% SDS gel and transferred to nitrocellulose membranes (Thermoscientific). Membranes were blocked with Odyssey Casein blocking buffer (LiCor) for one hour and incubated with mouse HNF4A (1:2500; ab41898, Abcam) and rabbit GAPDH (1:5000; G9545, Sigma) primary antibodies diluted in Odyssey Casein blocking buffer overnight. Following PBS-Tween washes, the membranes were incubated with anti-mouse IRDye680 (1:15,000; 926-32220, LiCor) and anti-rabbit IRDye800 (1:15,000; 926-32211, LiCor) diluted in Odyssey Casein blocking buffer for one hour. Membranes were scanned on the Odyssey^®^ CLx Infrared imaging system (LiCor) and band intensities were quantified using Image Studio Lite V5.2.

### Chromatin immunoprecipitation (ChIP)

The ChIP assay was performed as previously described (49) with minor modifications. Briefly, chromatin was extracted from 10 million HepG2 cells and was sonicated (Vibra Cell VCX130 Sonicator, Sonics) to yield DNA fragments of ~500 bp, as determined experimentally. Lysates were diluted 1:10 with immunoprecipitation (IP) buffer. The diluted chromatin was pre-cleared overnight using Dynabeads protein G (Thermofisher). Immunoprecipitation reactions were performed overnight at 4 °C using 10 μl anti-HNF4A (ab41898, Abcam) antibody conjugated to 50 μl of Dynabeads G.

DNA was recovered using Phenol:Chloroform:Isoamyl alcohol (Invitrogen) and precipitated with 0.1 volume of NaCl and 2.5 volumes of ethanol using Ambion Linear Acrylamide (5 mg/mL) as a carrier. qPCR was performed on 1 μl of pre-cleared or immunoprecipitated chromatin using the primers HNF4A motif F: 5’-CCCTGTCCTGACACTTGGTT-3’ and HNF4A motif R: 5’-CTCTGCATACCTTTGGAGGA-3’ to generate a 202 bp amplicon encompassing the target HNF4A motif. Control primers used for the amplification of a 77 bp positive binding site were ABCC6 promoter F: 5’AGCCCATTGCATAATCTTCTAAGT-3’ and ABCC6 promoter R: 5’-ATGGAGACCGCGTCACAG-3’, and a 110 bp negative binding site were β-globin F: 5’-AGGACAGGTACGGCTGTCATC-3’ and β-globin R: 5’-TTTATGCCCAGCCCTGGCTC-3’. Enrichment of each amplicon was determined after normalization to the pre-cleared input chromatin, after subtracting the ‘no antibody’ control.

For deep sequencing using the Illumina MiSeq, a two-step PCR approach was performed using the pre-cleared and immunoprecipitated chromatin as templates. In the first step, HNF4A motif primers modified to include an 18 bp adaptor-specific sequence were used to generate amplicons tagged at both ends by adaptor priming sites. The amplicons were purified using AMPure XP beads and were subjected to a second round of PCR using unique pairs of Illumina index primers. The indexed amplicons were purified using AMPure XP beads and quantified using the Qubit dsDNA HS Assay Kit. Amplicons were pooled to equal representation and 150 bp paired-end sequencing was performed. Quality control, clean-up of reads and alignment to the human reference genome (GRCh37/hg19) were performed using a web-based open source platform, Galaxy. Allelic frequencies were determined using the Integrative Genomics Viewer (50).

### Zebrafish assays

Wild type (WIK), transgenic and mutant fish lines were maintained according to established protocols (51). Single-cell WIK embryos were injected with 1 nl of 30 ng/µl pTol2Gata2 (46) DNA construct containing either of the alleles and 90 ng/µl Tol2 transposase mRNA (52). Injected embryos were screened for enhancer expression under the Leica M205 FA epifluorescence microscope. Fluorescence-positive embryos were raised under standard conditions to F1 adults. These were outcrossed and the resulting embryos were imaged for fluorescence at 4 dpf using the Leica M205 FA epifluorescence microscope (Leica Applications Suite) and Nikon C2 confocal microscope (Nikon NIS-Elements).

### Zebrafish histology

21 dpf larvae were fixed in 4% paraformaldehyde, embedded in paraffin and 3 micron sections were cut onto superfrost slides. Colorimetric *in situ* hybridization was carried out using a digoxigenin-labeled GFP riboprobe synthesized from a pCS2-EGFP plasmid using T7 RNA polymerase (Roche). Anti-DIG-alkaline phosphatase antibody (Roche) was used for detection, followed by visualization with nitro blue tetrazolium and 5-bromo-4-chloro-3-indolylphosphate (NBT/BCIP) (Roche). Sections were imaged using brightfield microscopy on a Nikon CS2 microscope and Nikon NIS-Elements software.

### Statistical analysis

GraphPad PRISM 7 was used for performing all statistical analysis. One-way ANOVAs, (Tukey’s multiple comparisons tests) were used for estimating the statistical significance of the dual luciferase assays and the ChIP-PCR assay. Unpaired *t*-tests were used for estimating the statistical significance of the siRNA-luciferase assays, quantitative PCR and western blotting data. All data are presented as the mean ± standard deviation (S.D.).

## ACKNOWLEDGEMENTS

This research was supported by Health Research Council of New Zealand grant #15/623 to T.R.M., J.M.O. and J.A.H. We thank Noel Jhinku for expert management of the Otago Zebrafish Facility, and Rob Day for assistance with the MiSeq analysis.

## CONFLICT OF INTEREST STATEMENT

The authors declare no conflict of interest.

